# Enhancing CRISPR prime editing by reducing misfolded pegRNA interactions

**DOI:** 10.1101/2023.08.14.553324

**Authors:** Weiting Zhang, Karl Petri, Junyan Ma, Hyunho Lee, Chia-Lun Tsai, J. Keith Joung, Jing-Ruey Joanna Yeh

## Abstract

CRISPR prime editing (**PE**) requires a Cas9 nickase-reverse transcriptase fusion protein (known as PE2) and a prime editing guide RNA (**pegRNA**), an extended version of a standard guide RNA (**gRNA**) that both specifies the intended target genomic sequence and encodes the desired genetic edit. Here we show that sequence complementarity between the 5’ and the 3’ regions of a pegRNA can negatively impact its ability to complex with Cas9, thereby potentially reducing PE efficiency. We demonstrate this limitation can be overcome by a simple pegRNA refolding procedure, which improved ribonucleoprotein-mediated PE efficiencies in zebrafish embryos by up to nearly 25-fold. Further gains in PE efficiencies of as much as 6-fold could also be achieved by introducing point mutations designed to disrupt internal interactions within the pegRNA. Our work defines simple strategies that can be implemented to improve the efficiency of PE.

---

PE is a versatile gene-editing technology that enables programmable installation of any nucleotide substitution and small insertions/deletions without requiring a DNA donor template or the introduction of DNA double-strand breaks^1^. It utilizes a Cas9 nickase (**nCas9**)-reverse transcriptase (**RT**) fusion protein called PE2 and a prime editing guide RNA (**pegRNA**, **Fig. 1a**). Like a gRNA, the pegRNA directs nCas9 to the target DNA site specified by the 5’ spacer sequence. The non-target DNA strand nicked by nCas9 then anneals to a complementary primer binding site (**PBS**) at the 3’ end of the pegRNA. Subsequently, the adjacent reverse transcriptase template (**RTT**) also encoded in the pegRNA is reverse transcribed by the PE2 RT generating a 3’ DNA “flap” that encodes the desired edit^2^. Although PE has been successfully employed in mammalian cells, plants, *Drosophila*, zebrafish and mice, the editing efficiencies observed are generally lower than those observed with other forms of CRISPR/Cas-based editing (e.g., nucleases and base editors)^2–6^.

**Fig. 1.**
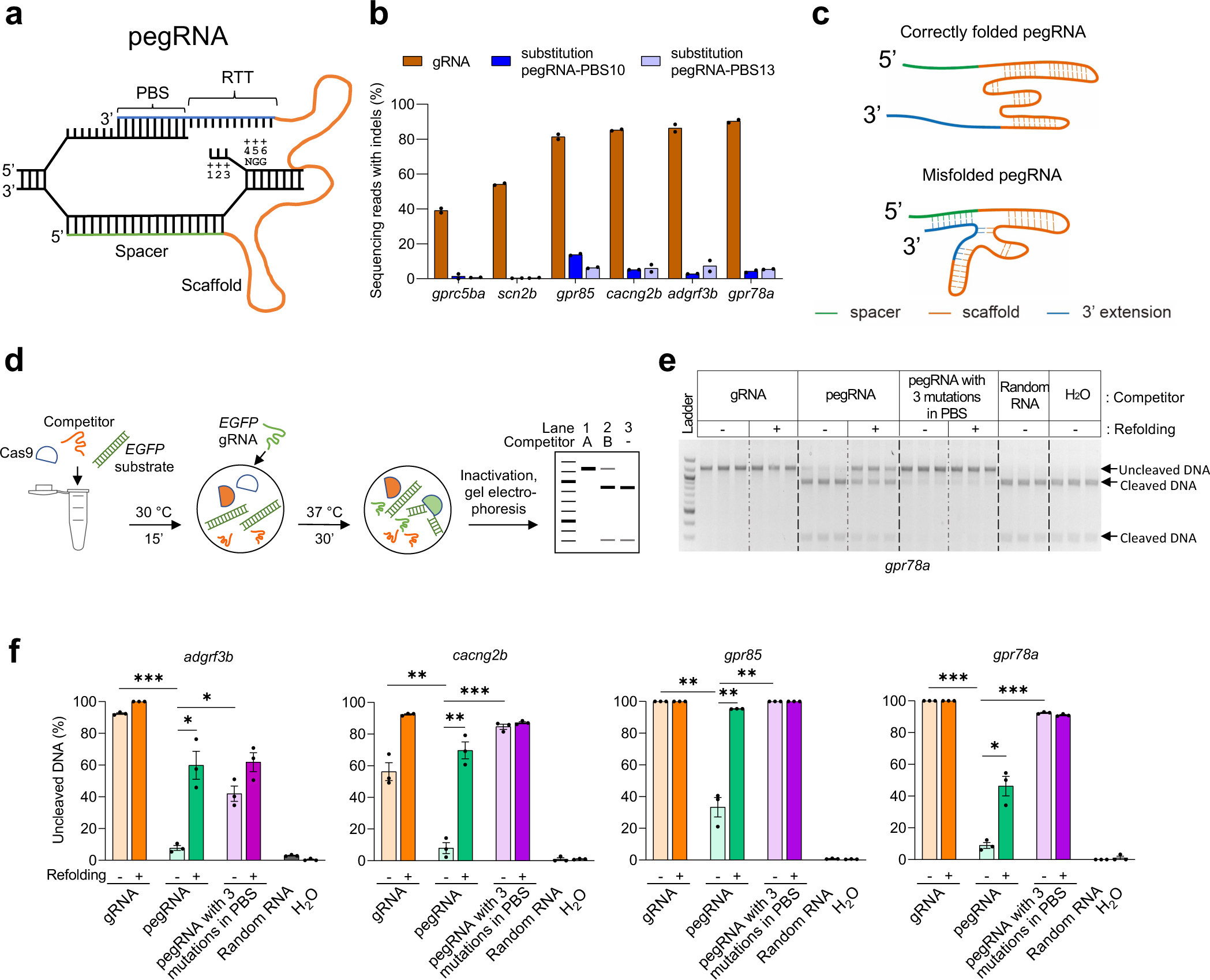
Improving *in vitro* SpCas9 binding efficiencies of pegRNA by refolding. **a**, Schematic illustrating the hybridization of a pegRNA and its target DNA. Four segments of a pegRNA are shown. PBS, Primer Binding Site; RTT, Reverse Transcriptase Template. Target DNA positions (as well as the corresponding sequences in the RTT) are numbered counting from the SpCas9-induced nick towards ‘NGG’, the protospacer adjacent motif (PAM). **b**, Mutation frequencies induced by SpCas9 with gRNAs and pegRNAs in zebrafish. All pegRNAs carried a single nucleotide substitution at position +5 or +6, with RTT lengths of 14- or 15-nucleotide (nt), and PBS lengths of 10-nt (pegRNA-PBS10) or 13-nt (pegRNA-PBS13). Target loci are indicated at the bottom. SpCas9 protein was complexed with gRNA or pegRNA at a molar ratio of 1:2 (0.6 µM of gRNA or pegRNA). **c**, Schematic illustrating hypothetical conformations of correctly folded and misfolded pegRNAs. The spacer is shown in green, Cas9-binding scaffold in orange and 3’ extension including PBS and RTT in blue. Dotted lines indicate potential base pairings. **d**, Schematic illustrating the *in vitro* competition assay for Cas9 binding and substrate cleavage. Possible outcomes of the assay are shown in a representative gel. Lane 1 shows the addition of Competitor A with a high SpCas9-binding affinity resulting in 100% inhibition of cleavage of DNA substrates (1.2 kilobase pairs). Lane 2 shows the addition of Competitor B with a low SpCas9-binding affinity yielding a mix of uncleaved and cleaved (900 and 300 base pairs) DNA substrate. Lane 3 shows the reaction without any competitor resulting in 100% cleaved DNA products. **e**, Agarose gel image showing the results of *in vitro* SpCas9 cleavage of DNA substrate in the presence of gRNA or pegRNAs targeting *gpr78a* as competitors, with or without refolding (indicated on top of the gel). Random RNA isolated from tolura yeast was used as a negative control. Assays were performed in triplicate. **f**, Percentage of uncleaved DNA substrate in the presence or absence of competitor gRNA or pegRNA calculated using data from Fig. 1e and **Supplementary Fig. 1**. Competitor gRNA and pegRNA target loci are indicated at the top. Competitor types are shown at the bottom. Dots represent individual data points, bars the mean and error bars ± s.e.m. Unpaired two-tailed *t*-test with equal variance was used to compare non-refolded gRNA vs non-refolded pegRNA, non-refolded pegRNA vs non-refolded pegRNA with 3 mutations in PBS, and non-refolded vs refolded pegRNAs. \**p* < 0.05, \*\**p* < 0.01, \*\*\**p* < 0.001.

To explore potential mechanisms for the lower editing frequencies observed with PE, we tested whether the 3’ PBS/RTT segment might impact the ability of a pegRNA to function with standard SpCas9 nuclease. To do this, we compared the mutation frequencies induced by SpCas9 with matched pairs of pegRNAs and standard gRNAs (i.e., lacking the PBS and RTT 3’ extensions) targeting the same spacer sequences. We performed these comparisons in 1-cell stage zebrafish embryos by injecting Cas9 protein complexed with equimolar concentrations of either pegRNAs or gRNAs targeting six different endogenous gene loci. To quantify indel frequencies induced at these sites, we extracted genomic DNA from embryos one day following injection and performed targeted amplicon next-generation sequencing (**NGS**). We found that across all six target spacer sites, lower indel frequencies were observed with pegRNAs than with their matched gRNAs (**Fig. 1b**), suggesting that the presence of the 3’ extension in a pegRNA decreases Cas9-induced gene editing.

One potential explanation for the lower Cas9 activities we observed with pegRNAs compared with standard gRNAs is that interactions between complementary sequences in the 5’ spacer and 3’ PBS and RTT might cause misfolding of pegRNAs in a way that decreases their abilities to complex with Cas9 protein (**Fig. 1c**). This possibility seemed likely given that previous work has shown that even shorter length internal interactions between bases in standard gRNAs can stabilize alternative non-functional folding and low SpCas9-induced mutation efficiencies^7^. To test our hypothesis, we used a previously described *in vitro* assay^7^ that assesses the abilities of various gRNAs (and, in this case, pegRNAs) to complex with Cas9 protein. In this assay, various matched pegRNAs and gRNAs targeted to endogenous zebrafish gene spacer targets compete with a gRNA targeting an *EGFP* reporter gene sequence for complexation with Cas9 nuclease (**Fig. 1d**). The degree of successful competition by a given gRNA or pegRNA can be assessed by measuring cleavage of an *EGFP* target DNA site substrate included in each reaction (**Fig. 1d**). Using this assay, we compared matched gRNAs and pegRNAs targeted to four different genomic DNA sites, *gpr78a, adgrf3b*, *cacng2b* and *gpr85* and found that in all four cases the gRNAs could efficiently compete with the *EGFP*-targeted gRNA for binding to Cas9 (as judged by reduced cleavage of the *EGFP* DNA target site template), whereas the pegRNAs were substantially and significantly reduced in this capability (**Figs. 1e - 1f; Supplementary Fig. 1**). To test whether this reduced Cas9-binding capability of the pegRNAs might be caused in part by their 5’ vs 3’ complementarity, we performed additional *in vitro* experiments with matched pegRNAs in which we introduced three point mutations in the 3’ PBS region of the pegRNAs (**Supplementary Table. 1**) and found that these mutated pegRNAs showed significant increases in their competitive Cas9-binding activities (**Figs. 1e - 1f; Supplementary Fig. 1**).

Having determined that pegRNAs may suffer from reduced binding to Cas9 protein, we next considered whether refolding the pegRNAs could improve their Cas9 binding capabilities. We considered this hypothesis because it has been previously shown that refolding low activity gRNAs believed to have alternative folding configurations can significantly improve their abilities to mediate Cas9-induced indels^7^. We found that subjecting pegRNAs to a refolding procedure consisting of heat denaturation followed by slow cooling (**Methods**) significantly improved their abilities to bind Cas9 in the *in vitro* DNA cleavage competition assay (although not to levels observed with matched standard gRNAs) (**Figs. 1e - 1f; Supplementary Fig. 1**). Consistent with this, we also observed that re-folding of pegRNAs prior to formation and injection of pegRNA-Cas9 ribonucleoprotein complexes into zebrafish embryos also increased Cas9-mediated indel frequencies by up to 2.8-fold for four of the seven target sites tested (**Supplementary Fig. 2**).

Next, we evaluated whether refolding the pegRNAs would also increase PE efficiencies in zebrafish embryos. For this experiment we used 17 pegRNAs from our earlier study^4^ that were designed to introduce various types of mutations (base substitutions, insertions, or deletions) and that had exhibited a range of PE efficiencies from barely active to the best performers. We found that refolding prior to embryo injection significantly increased the frequencies of pure PE (defined as the alleles containing only the pegRNA-specified edits without any other mutations) frequencies for nine of these 17 pegRNAs (**Figs. 2a – 2b**). Significant increases were observed with seven of 12 pegRNAs designed to introduce base substitutions (increases ranging from 2.6- to 24.7-fold) (**Fig. 2a**) and 2 of 3 pegRNAs designed to create insertion pegRNAs (increases ranging from 1.7- to 4.6-fold) (**Fig. 2b**). We did not observe significant increases in PE efficiencies with the two pegRNAs designed to induce deletions (**Fig. 2b**), perhaps due relatively lower degree of complementarity between the 5’ spacer and 3’ RTT because of the deletion encoded in the latter region. In general, increases in pure PE frequencies due to re-folding were accompanied by significant increases in non-pure PE frequencies (**Supplementary Figs. 3a – 3b**) and therefore PE purity values (defined as the percentage of pure PE edits out of total edits) did not show significant differences for most of the pegRNAs we assessed (**Supplementary Figs. 3c – 3d**). The only exceptions were two *scn2b* substitution pegRNAs that actually showed significantly improved PE purities with re-folding and the *cacng2b* deletion pegRNA that showed significantly reduced PE purity with refolding (**Supplementary Figs. 3c – 3d**).

**Fig. 2.**
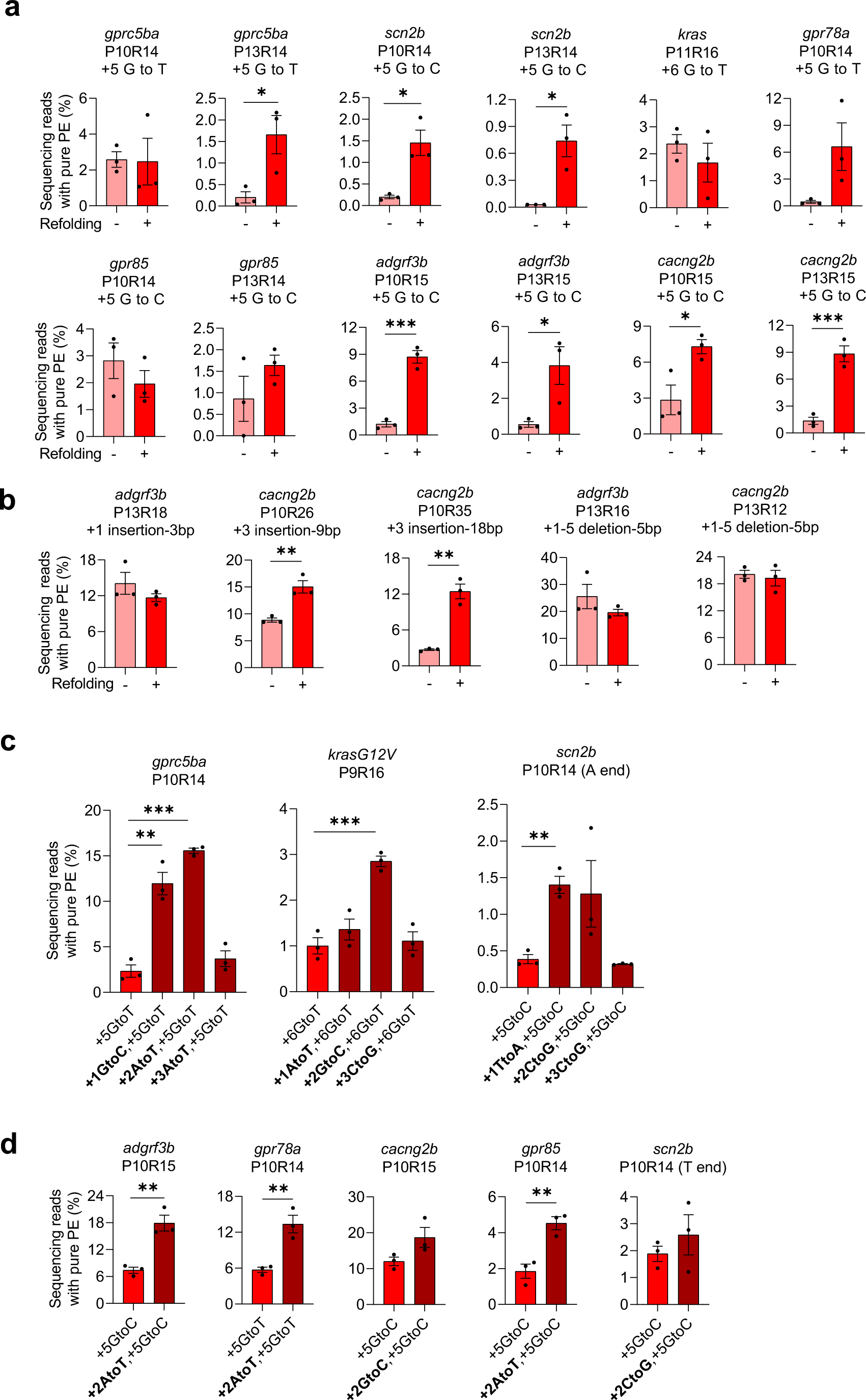
Improving prime editing efficiencies in zebrafish by pegRNA refolding and mutations in RTT. **a-b**, Pure PE frequencies of non-refolded and refolded substitution pegRNAs (**a**) and insertion or deletion pegRNAs (**b**) with PE2 in zebrafish. Target loci, PBS lengths (labeled as “P” followed by the number of nucleotides), RTT lengths (labeled as “R” followed by the number of nucleotides), and pegRNA-specified edits (denoted as the position of the edit followed by the edit) are shown at the top. Pure PE represents sequencing reads containing only the pegRNA-specified mutations. **c-d**, Pure PE frequencies with refolded pegRNAs carrying additional RTT mutations (at +1, +2 or +3) and PE2 in zebrafish. Target loci, PBS, and RTT lengths are shown at the top and pegRNA-specified edits are shown at the bottom. All pegRNAs had 3 or 4 thymine (T) nucleotides at the 3’ end except for the ones labeled ‘A end’ for *scn2b* in which the terminal Ts were replaced with adenine (A) nucleotides. Dots represent individual data points (*n* = 3 biologically independent replicates, 5-10 embryos per replicate), bars the mean and error bars ± s.e.m. \**p* < 0.05, \*\**p* < 0.01, \*\*\**p* < 0.001 (unpaired two-tailed *t*-test with equal variance).

Lastly, we tested whether introduction of single point mutations in the RTT region to reduce complementarity of pegRNA 5’ and 3’ regions might also lead to increases in PE efficiencies. Specifically, we explored the effects of creating mutations at the +1, +2, or +3 positions of the RTT (+1 defined as the first nucleotide just 5’ to the first nucleotide of the PBS and +2 and +3 being further upstream within the pegRNA, **Fig. 1a**) on PE efficiencies in zebrafish embryos. To do this, we introduced RTT +1, +2, or +3 mutations into three pegRNAs that each mediated low PE frequencies even after re-folding (0.39% to 2.33%; **Fig. 2c**) and that each encoded a single nucleotide substitution edit (encoded at RTT positions +5 or +6). Following re-folding, complexation with PE2, and injection into zebrafish embryos, we found that pegRNAs harboring mutations at the RTT +1 and +2 positions could make PE more efficient than their unmutated pegRNA counterparts (**Fig. 2c; Supplementary Fig. 4a**). Based on these results, we introduced a mutation at the RTT +2 position in five additional pegRNAs specifying a single edit and found that the mutation increased mean pure PE frequencies in all five cases (**Fig. 2d**). Three of the five mutated pegRNAs showed statistically significant increases in pure PE frequencies and two of these three pegRNAs also showed significant increases in non-pure PE frequencies (**Fig. 2d; Supplementary Fig. 4b**). Overall, a mutation at the RTT +2 position increased pure PE frequency up to 6.7-fold (mean 2.4-fold) (**Figs. 2c – 2d**). PE purity was unchanged except for one pegRNA (*cacng2b*) that showed a modest but statistically significant increase (**Supplementary Fig. 4c**). Testing of three re-folded pegRNAs harboring the RTT +2 mutations showed that they all induced higher indel frequencies with SpCas9 nuclease in zebrafish embryos than matched re-folded pegRNAs without the RTT +2 mutations (**Supplementary Fig. 5**), suggesting that the +2 mutations function to enhance the abilities of these pegRNAs to form functional complexes with Cas9 protein.

Our work delineates and validates two simple and general strategies for improving the efficiency of PE that can be readily practiced by any investigator. Delivery of PE components as RNP complexes offers multiple potential advantages relative to DNA or RNA delivery (e.g., increased efficiency, reduced off-target effects, and avoiding risk of integration events)^8–14^ and we show that combining pegRNA re-folding with the introduction of point mutations at the RTT +1 or +2 position can increase RNP-induced pure PE frequencies by as much as 29-fold in zebrafish embryos (**Fig. 1**, **Fig. 2** and **Supplementary Table. 2**). Our previous results showing that RNP-mediated PE functions in both zebrafish embryos and cultured human cells^4^ suggest that these simple strategies should likely improve PE in other settings such as human and other cell types as well. The introduction of RTT mutations to reduce pegRNA 5’ and 3’ complementarity can also be used when practicing PE technology using non-RNP-delivery methods such as DNA or RNA transfection, transduction, and/or injection. Notably, Li et al. have reported that introducing additional mutations in the RTT +1 to +3 positions can also enhance PE efficiency in human cells via DNA transfection^15^. One important consideration in using the RTT mutation strategy is to ensure that the additional change introduced is either silent (if in a coding region) or otherwise benign in its effect^15^. Corroborating our findings, Ponnienselvan et al. recently reported that 3’ truncated pegRNAs are preferentially loaded onto Cas9 and the prime editor protein, potentially due to reduced pegRNA 5’-3’ interactions^14^. Although beyond the scope of this current work, it will be interesting to explore whether pegRNA re-folding and/or RTT mutation might be combined with other previously described strategies for improving PE efficiencies (e.g., adding various structured RNA motifs to the 3’ termini of pegRNAs^16, 17^ or PE proteins with improved architectures and activities^16, 18, 19^).

Our findings also have more broad implications for both the design of pegRNAs and the range of genomic spacer sequences that can be targeted by PE for recognition and editing. Previous work has shown that the length and base composition of pegRNA PBS and RTT sequences can influence PE efficiency for any given target spacer site^14, 20–22^. Our work adds another parameter (internal complementarity between the 5’ spacer and 3’ PBS/RTT regions of the pegRNA) to be considered as one designs and tests various combinations to optimize PE activity. Accounting for this additional consideration may impact the nature of spacer sequences that can efficiently be targeted (e.g., higher GC content and/or melting temperature may actually be undesirable) and the length and composition of PBS/RTT sequences that can be used. We envision that the generation of larger datasets of optimized pegRNAs and consideration of all the parameters that can influence pegRNA activities (including internal complementarity) may yield improved rules and software for in silico design in the future.

## Acknowledgments

This work was supported by the Hassenfeld Scholar Award (to J.-R.J.Y.), NIH R01 GM134069 (to J.-R.J.Y.), and NIH RM1 HG009490 (to J.K.J.). K.P. was funded by the Deutsche Forschungsgemeinschaft (DFG, German Research Foundation) – Projektnummer 417577129. J.M. received support from the China Scholarship Council (CSC, 201808210354). We thank L. Paul-Pottenplackel for help with revising the manuscript.

## Author Contributions

W.Z., K.P., J.K.J., and J.-R.J.Y. designed the project; W.Z., K.P., and H.L. performed the experiments; W.Z. and J.-R.J.Y. developed the methods; W.Z., K.P., and H.L. performed informatic analysis; J.M. and C.-L.T. provided resources; J.K.J. and J.-R.J.Y. provided oversight; W.Z., J.K.J., and J.-R.J.Y. wrote the manuscript with input from all the authors.

## Supplementary Figure Legends

**Supplementary Fig. 1.**
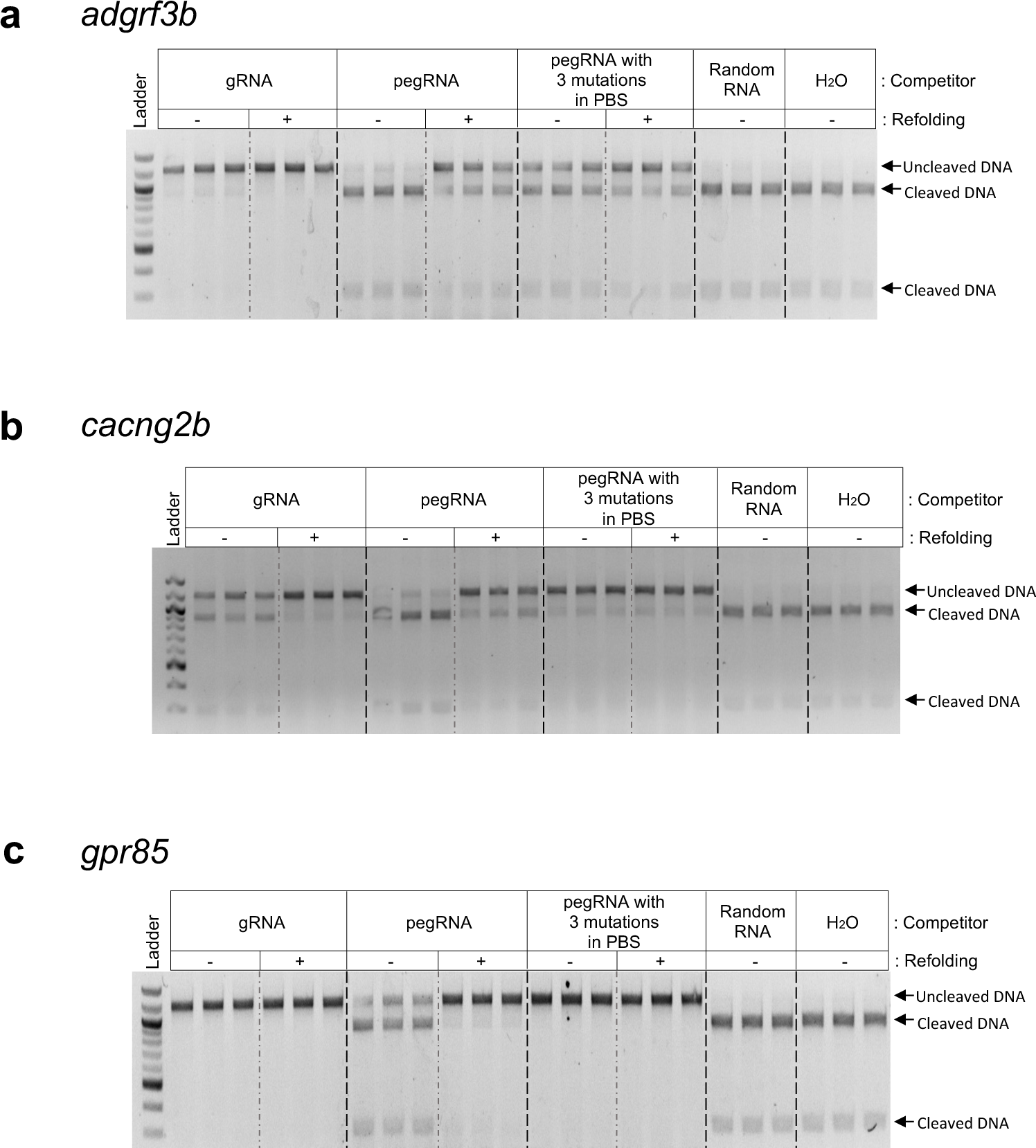
*In vitro* competition and DNA cleavage assay to assess the SpCas9-binding affinity of gRNAs and pegRNAs. **a-c**, Agarose gel showing *in vitro* SpCas9 cleavage of *EGFP* DNA substrate in the presence of gRNAs and pegRNAs targeting *adgrf3b* (**a**), *cacng2b* (**b**), or *gpr85* (**c**) as competitors with and without refolding. The substitution pegRNAs with three mutations in PBS had mutations that could weaken potential interactions between 5’ spacer and 3’ PBS (**Supplementary Table 1**). Random RNA isolated from tolura yeast was used as a negative control. The assays were performed in triplicate. Percentage of uncleaved DNA substrate in the presence or absence of competitor gRNA or pegRNA calculated using this data are shown in Fig. 1f.

**Supplementary Fig. 2.**
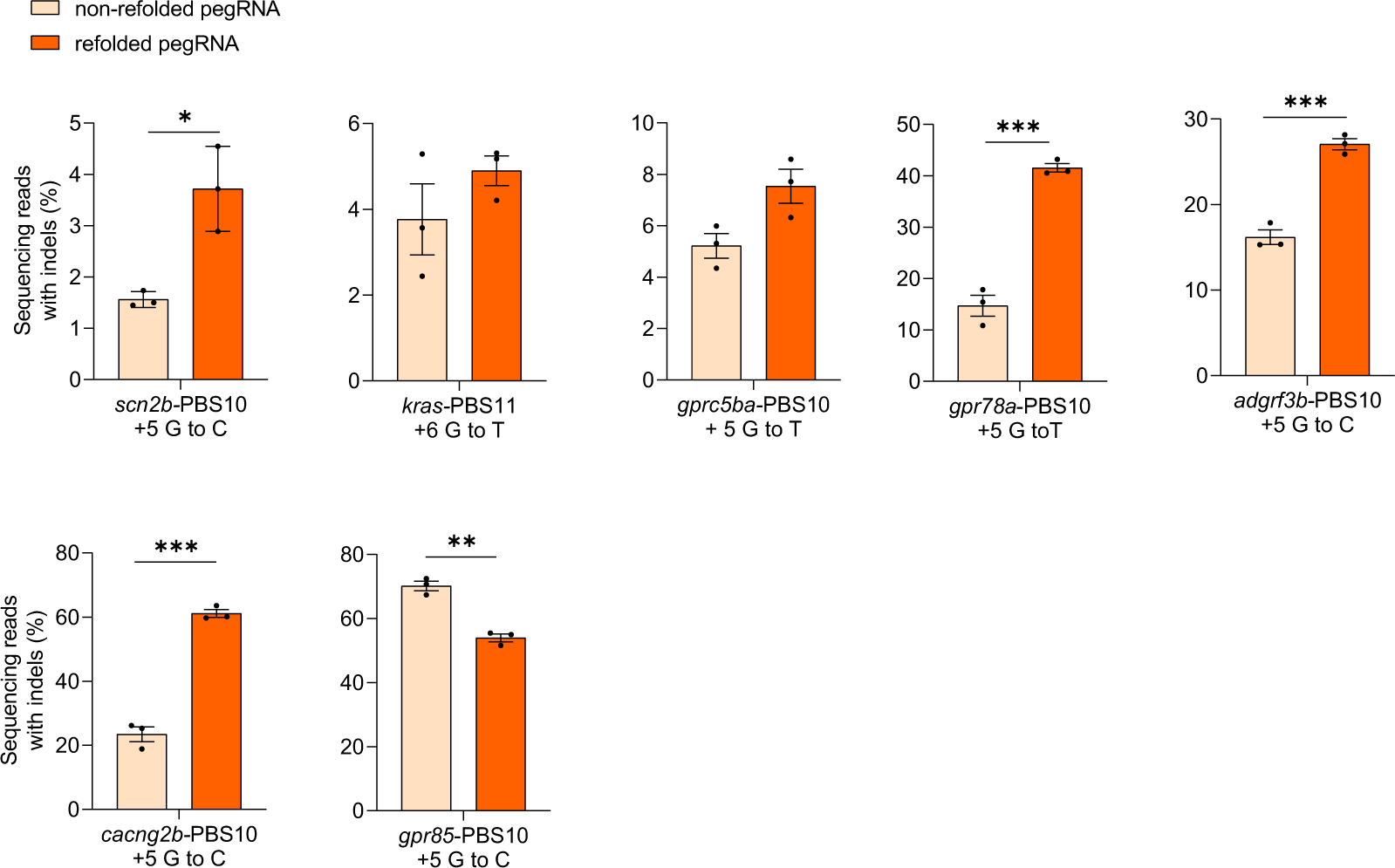
Indel frequencies of non-refolded and refolded pegRNAs with SpCas9 in zebrafish. Indel frequencies in zebrafish induced by SpCas9 complexed with non-refolded or refolded pegRNAs at a molar ratio of 1:2 (1.8 µM of pegRNA). Target loci, PBS lengths, and pegRNA-specified edits are indicated at the bottom. Dots represent individual data points (*n* = 3 biologically independent replicates, 5-10 embryos per replicate), bars the mean and error bars ± s.e.m. Results of unpaired two-tailed *t*-test with equal variance are shown in \**p* < 0.05, \*\**p* < 0.01,\*\*\**p* < 0.001.

**Supplementary Fig. 3.**
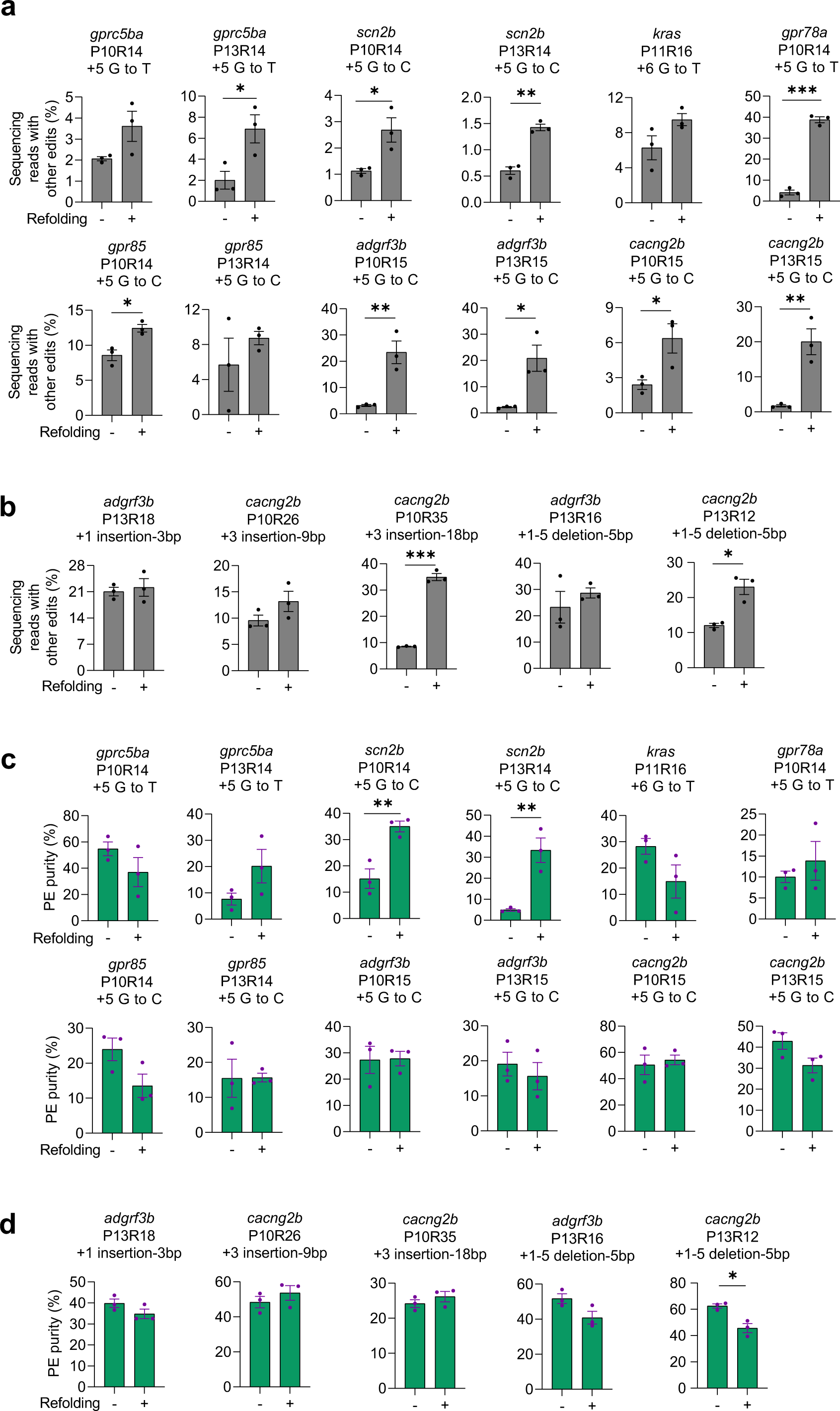
Purities of prime editing with PE2 and non-refolded or refolded pegRNAs in zebrafish. a-b. Frequencies of non-pure PE edits (labeled as other edits) mediated by PE2 with non-refolded and refolded substitution pegRNAs (**a**) and insertion or deletion pegRNAs (**b**). **c-d.** PE purities (%) calculated as pure PE over total edit (pure and non-pure PE) frequencies for substitution pegRNAs (**c**) and insertion or deletion pegRNAs (**d**). Target loci, PBS lengths (labeled as “P” followed by the number of nucleotides), RTT lengths (labeled as “R” followed by the number of nucleotides), and pegRNA-specified edits are indicated at the top. Dots represent individual data points (*n* = 3 biologically independent replicates, 5-10 embryos per replicate), bars the mean and error bars ±s.e.m. Results of unpaired two-tailed *t*-test with equal variance are shown in \**p* < 0.05, \*\**p* < 0.01,\*\*\**p* < 0.001.

**Supplementary Fig. 4.**
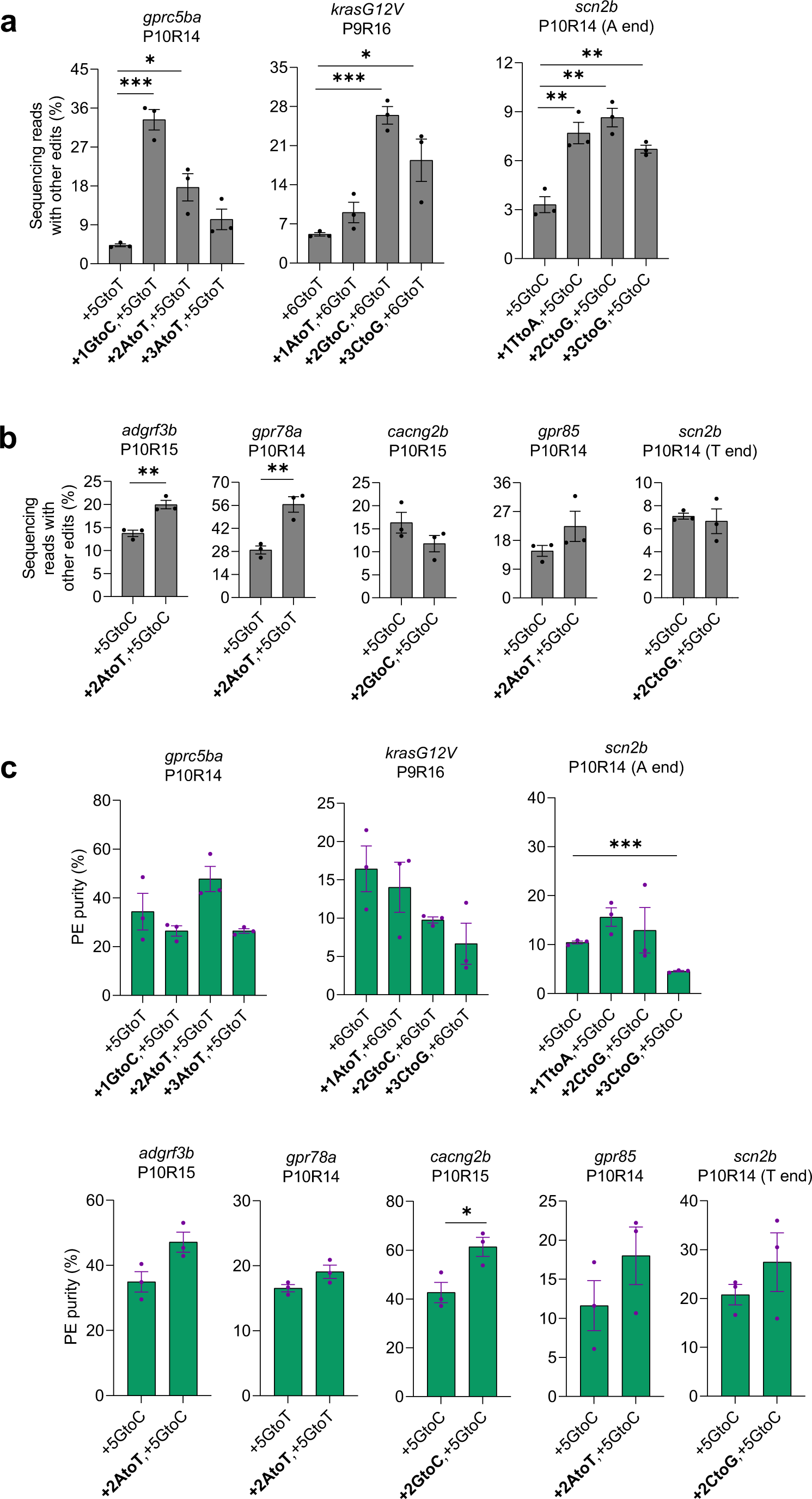
Purities of prime editing with PE2 and refolded pegRNAs carrying additional RTT mutations in zebrafish. **a-b.** Frequencies of non-pure PE edits (labeled as other edits) mediated by PE2 with refolded pegRNAs with or without an additional mutation in the RTT (at +1, +2 or +3). **c**. PE purities calculated as pure PE over total edit (pure and non-pure PE) frequencies. Target loci, PBS lengths (labeled as “P” followed by the number of nucleotides), and RTT lengths (labeled as “R” followed by the number of nucleotides) are shown at the top. pegRNA-specified edits (denoted as the position of the edit followed by the edit) are shown at the bottom. Dots represent individual data points (*n* = 3 biologically independent replicates, 5-10 embryos per replicate), bars the mean, error bars ± s.e.m. Results of unpaired two-tailed *t*-test with equal variance are shown in \**p* < 0.05, \*\**p* < 0.01,\*\*\**p* < 0.001.

**Supplementary Fig. 5.**
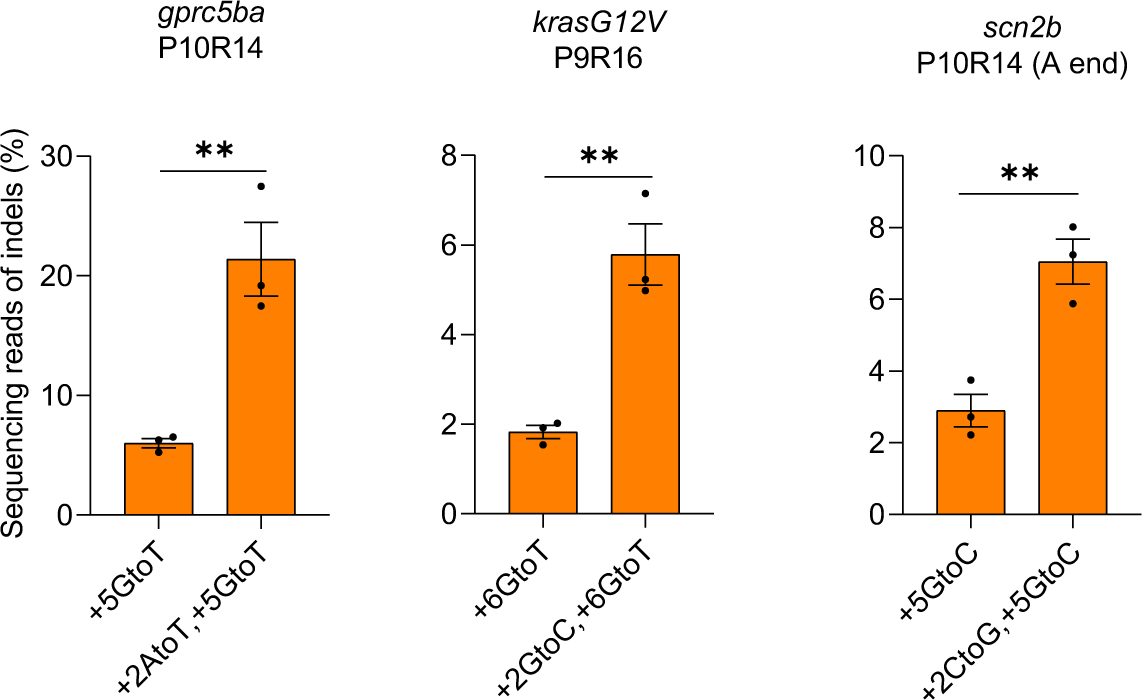
Indel frequencies in zebrafish induced by SpCas9 complexed with refolded pegRNAs with or without RTT mutation at +2 position. SpCas9 protein and pegRNA were combined at a molar ratio of 1:2. (1.8 µM of pegRNA). Target loci, PBS lengths (labeled as “P” followed by the number of nucleotides), and RTT lengths (labeled as “R” followed by the number of nucleotides) are shown at the top. pegRNA-specified edits (denoted as the position of the edit followed by the edit) are shown at the bottom. Dots represent individual data points (*n* = 3 biologically independent replicates, 5-10 embryos per replicate), bars the mean and error bars ±s.e.m. Results of unpaired two-tailed *t*-test with equal variance are shown in \*\**p* < 0.01.

## Methods

### Production of gRNAs and pegRNAs

The gRNAs and pegRNAs (Supplementary Table 3) used in this work were synthesized by *in vitro* transcription. The DNA templates for *in vitro* transcription were constructed by one-step PCR, using a C9E constant oligonucleotide containing the enhanced SpCas9 scaffold^4^, forward primer carrying the SP6 promoter and target-specific spacer, and the reverse primer with the 3’ extension containing the primer binding site and RTT. Primer sequences and PCR formulation are listed in Supplementary Table 4. PCR reactions were conducted with Phusion High-Fidelity DNA polymerase (New England Biolabs, no. M0530L) using the following cycling program: 98 °C for 30 s, followed by 35 cycles of 98 °C for 10 s, 65 °C for 30 s, and 72 °C for 15 s, followed by a final 72 °C extension for 5 min. The PCR products were purified using the Monarch PCR & DNA Cleanup Kit (New England Biolabs, no. T1030L). *In vitro* transcription of pegRNAs and gRNAs was performed using the HiScribe SP6 RNA Synthesis Kit (New England Biolabs, no. E2070S), purified using the Monarch RNA Cleanup Kit (New England Biolabs, no. T2030L) and eluted in water.

### Purification of PE2 and SpCas9 protein

PE2 protein was purified as previously described^4^. SpCas9 protein was purified as described in ^23^ with some modifications. Chemically competent Rosetta (DE3) competent cells (Novagen, no. 70954) were transformed with pET-28b-Cas9-His (Addgene, no. 47327) by heat shock following the manufacturers’ instructions. 25 milliliters of overnight culture grown from a single colony in Luria Bertani (LB) medium with 50 μg ml^−1^ kanamycin was transferred into 670 ml of autoinduction medium (24 g l^−1^ yeast extract, 12 g l^−1^ soy peptone, 12.5 g l^−1^ potassium phosphate dibasic, 2.3 g l^−1^ potassium phosphate monobasic, 0.4% glycerol) containing 50 μg ml^−1^ kanamycin. The culture was incubated at 37°C for about 4 hours until the OD_600_ of the culture had reached 1.0 - 2.0 and was then switched to 18 °C for another 24 hours. Cells were collected and resuspended in lysis buffer consisting of 20 mM Tris-HCl pH 8.0, 20 mM imidazole, 150 mM NaCl, 0.1% Triton X-100, 5 mM 2-mercaptoethanol and cOmplete Protease Inhibitor Cocktail (Roche, no. 11697498001). The cell suspension was subjected to sonication (Qsonica) for 2 minutes (20-second pulses and 20-second rest between pulses) at 4°C. Lysate was cleared by centrifugation at 14,000 rpm for 10 min at 4 °C. The supernatant was mixed with 2 ml Ni-NTA agarose (Qiagen, no. 30250) and kept on a rotator at room temperature for 1 h.

Subsequently, the supernatant-agarose mixture was loaded onto an Econo-Column chromatography column (Bio-Rad, no. 7374011) and the supernatant flowed through by gravity. The flow-through was re-loaded onto the same column once more. The column was then washed with 80 ml of wash buffer (20 mM Tris pH 8, 20 mM Imidazole, 500 mM NaCl). Protein was eluted with 20 mM Tris pH 8, 250 mM Imidazole, and 300 mM NaCl, analyzed for purity by SDS–polyacrylamide gel electrophoresis (PAGE), and the elution buffer was replaced with 20 mM Tris pH 8.0, 200 mM KCl, 10 mM MgCl_2,_ and 10% glycerol by dialysis using Slide-A-Lyzer G2 Dialysis Cassettes with 20-*kDa* cutoff (Thermo Fisher Scientific, no. 87737) at 4 °C overnight. Protein concentration was determined with UV absorbance at 280 nm on a NanoDrop spectrometer, and purified proteins were stored at −80 °C.

### Refolding of pegRNAs and RNP complexation of refolded pegRNAs with Cas9 or PE2

PegRNAs in water were refolded by heating at 98°C for 2 min and slowly cooling down at a rate of −0.1°C per second to 30 °C. Refolded pegRNAs were immediately added to the SpCas9 or PE2 protein, mixed gently, spun for 10 sec at 6000 RPM in a mini centrifuge (Fisherbrand™), and followed by a 10 min incubation at 30 °C.

### *In vitro* competition assay for Cas9 binding and substrate cleavage

We tested the ability of pegRNAs and matched gRNAs to inhibit Cas9-induced cleavage of a DNA substrate (*EGFP*) by competing with the cognate *EGFP* gRNA for binding to Cas9^7^. For this assay 6 μl of Cas9 protein (16.67 ng/µl) in 400 mM KCl, 20 mM MgCl_2_, and 40 mM Tris-HCL at pH 8.0. was mixed with 2 μl of gRNA or pegRNA being tested, or random RNA (SIGMA, no R6625) in a molar ratio of 1:2, and 2 μl of 25 ng/μl *EGFP* DNA substrate (1200 bp, amplified using the primers listed in Supplementary Table 5). After incubating the mixture at 30 °C for 15 min, 2 μl of *EGFP* gRNA at the same molar concentration as the test gRNAs and pegRNAs, was added to the mixture and the cleavage reaction performed at 37 °C. After 30 min, the reactions were stopped by adding a gel loading dye (New England Biolabs, no. B7024A) followed by heat inactivation at 80 °C for 10 min. The cleaved DNA samples were separated on a 2% agarose gel by electrophoresis. The fluorescent intensity of each band was calculated by dividing its total fluorescent intensity measured in Image J/FIJI Gels by its band size in bp, yielding a unit fluorescent intensity for each band. The cleavage percentages were calculated by dividing the unit fluorescent intensity of the cleaved band at 900 bp by the sum of the values for the non-cleaved band at 1200 bp and the cleaved band at 900 bp.

### Zebrafish husbandry

All zebrafish husbandry and experiments were approved by the Massachusetts General Hospital Subcommittee on Research Animal Care and performed under the guidelines of the Institutional Animal Care and Use Committee at the Massachusetts General Hospital.

### Zebrafish gene editing with Cas9 and PE2

Microinjections were performed using the one-cell stage of TuAB zebrafish embryos. Each embryo was injected with 2 nl of the RNP mixture at a Cas9 or PE2 protein to gRNA or pegRNA molar ratio of 1:2 (750 ng/µl of PE2 with 240 ng/µl of pegRNA, Fig. 2; 500 ng/µl of Cas9 with 3 mixed pegRNAs of 80 ng/µl each, Supplementary Figs. 2 and 5) and immediately transferred to an incubator at 32 °C for PE2 and 28.5 °C for Cas9. For Fig. 1b, six gRNAs, pegRNAs with a 10-nt PBS (pegRNA-PBS10), or pegRNAs with a 13-nt PBS (pegRNA-PBS13) were pooled together and mixed with Cas9 (20 ng/µl of each gRNA and 25 ng/µl of each pegRNA; Cas9 to gRNA/pegRNA molar ratio of 1:2) and injected as described above. One day post-fertilization, between five and ten embryos that developed normally from each condition were pooled and lysed in 10 mM Tris-HCl pH 8.0, 2 mM EDTA pH 8.0, 0.2% Triton X-100 and 100 μg ml^−1^ Proteinase K (5–6 μl of lysis buffer/embryo). Lysates were incubated at 50 °C overnight with occasional mixing, heated at 95 °C for 10 min to inactive Proteinase K, and stored at 4 °C.

### Targeted deep sequencing

Amplicons for targeted sequencing were generated in two PCR steps. In the first step (PCR1), regions containing the target sites were amplified from 1 μl of the zebrafish embryo lysate, using touchdown PCR with Phusion High-Fidelity Polymerase (NEB, no. M0530S) and primers containing partial sequencing adapters (Supplementary Table 6). For some samples, the products of PCR1 were purified using the Monarch PCR & DNA Cleanup Kit (New England Biolabs, no. T1030L) and deep sequenced at the MGH DNA Core. For the rest of the samples, the products of PCR1 were diluted 200-fold with water and used in the second PCR step (PCR2), where Illumina barcodes and P5/P7 sequences were attached to PCR1 products. The PCR2 product yield was assessed by agarose gel electrophoresis and pooled together in equal amounts. The PCR2 product pools were subjected to three steps of purification with the Monarch PCR & DNA Cleanup Kit (New England Biolabs, no. T1030L), Gel DNA Recover Kit (Zymoclean, no.11-300C), and paramagnetic beads (1:1 beads/sample) using the same purification protocol as with AMPure XP beads (Beckman Coulter, no. B37419AB). The product purity was assessed via capillary electrophoresis on a QIAxcel instrument (Qiagen) and quantified by spectrophotometry (NanoDrop®). The resulting sequencing libraries were sequenced using the MiSeq system (Illumina v.2 kit, 2 × 150 bp).

### Deep sequencing analysis

Sequencing data were analyzed with CRISPResso2^24^. Cas9 nuclease-treated samples were analyzed using Cas9 mode with a 5-bp quantification window size. PE-treated samples were analyzed using Prime editor mode with a 5-bp quantification window size and a 5-bp pegRNA extension quantification window size. CRISPResso2 was run with quality filtering (only those reads with an average quality score ≥30 were considered).

### Statistical analysis

For all bar graphs, mean and s.e.m. (only for samples with n > 2) were calculated and plotted using GraphPad Prism v.8. Statistical analysis of the significance level was conducted using an unpaired two-tailed t-test with equal variance in Microsoft Excel version 2301.

## Data availability

Deep sequencing data will be deposited in the NCBI Sequence Read Archive (project no: PRJNA995387).

## References

1. Anzalone, A.V., Koblan, L.W. & Liu, D.R. Genome editing with CRISPR-Cas nucleases, base editors, transposases and prime editors. Nat Biotechnol 38, 824–844 (2020).

2. Anzalone, A.V. et al. Search-and-replace genome editing without double-strand breaks or donor DNA. Nature 576, 149–157 (2019).

3. Bosch, J.A., Birchak, G. & Perrimon, N. Precise genome engineering in Drosophila using prime editing. Proc Natl Acad Sci U S A 118, e2021996118 (2021).

4. Petri, K. et al. CRISPR prime editing with ribonucleoprotein complexes in zebrafish and primary human cells. Nat Biotechnol 40, 189–193 (2022).

5. Lin, Q. et al. Prime genome editing in rice and wheat. Nat Biotechnol 38, 582–585 (2020).

6. Liu, Y. et al. Efficient generation of mouse models with the prime editing system. Cell Discov 6, 27 (2020).

7. Thyme, S.B., Akhmetova, L., Montague, T.G., Valen, E. & Schier, A.F. Internal guide RNA interactions interfere with Cas9-mediated cleavage. Nat Commun 7, 11750 (2016).

8. Kim, S., Kim, D., Cho, S.W., Kim, J. & Kim, J.S. Highly efficient RNA-guided genome editing in human cells via delivery of purified Cas9 ribonucleoproteins. Genome research 24, 1012–1019 (2014).

9. Raguram, A., Banskota, S. & Liu, D.R. Therapeutic in vivo delivery of gene editing agents. Cell 185, 2806–2827 (2022).

10. Burger, A. et al. Maximizing mutagenesis with solubilized CRISPR-Cas9 ribonucleoprotein complexes. Development 143, 2025–2037 (2016).

11. Kanchiswamy, C.N. DNA-free genome editing methods for targeted crop improvement. Plant Cell Rep 35, 1469–1474 (2016).

12. Svitashev, S., Schwartz, C., Lenderts, B., Young, J.K. & Mark Cigan, A. Genome editing in maize directed by CRISPR-Cas9 ribonucleoprotein complexes. Nat Commun 7, 13274 (2016).

13. Wang, D., Zhang, F. & Gao, G. CRISPR-Based Therapeutic Genome Editing: Strategies and In Vivo Delivery by AAV Vectors. Cell 181, 136–150 (2020).

14. Ponnienselvan, K. et al. Reducing the inherent auto-inhibitory interaction within the pegRNA enhances prime editing efficiency. Nucleic Acids Res (2023).

15. Li, X. et al. Highly efficient prime editing by introducing same-sense mutations in pegRNA or stabilizing its structure. Nat Commun 13, 1669 (2022).

16. Nelson, J.W. et al. Engineered pegRNAs improve prime editing efficiency. Nat Biotechnol 40, 402–410 (2022).

17. Zhang, G. et al. Enhancement of prime editing via xrRNA motif-joined pegRNA. Nat Commun 13, 1856 (2022).

18. Doman, J.L., Sousa, A.A., Randolph, P.B., Chen, P.J. & Liu, D.R. Designing and executing prime editing experiments in mammalian cells. Nat Protoc 17, 2431–2468 (2022).

19. Liu, P., et al. Improved prime editors enable pathogenic allele correction and cancer modelling in adult mice. Nat Commun 12, 2121 (2021).

20. Kim, H.K. et al. Predicting the efficiency of prime editing guide RNAs in human cells. Nat Biotechnol 39, 198–206 (2021).

21. Li, Y., Chen, J., Tsai, S.Q. & Cheng, Y. Easy-Prime: a machine learning-based prime editor design tool. Genome Biol 22, 235 (2021).

22. Lin, Q. et al. High-efficiency prime editing with optimized, paired pegRNAs in plants. Nat Biotechnol 39, 923–927 (2021).

## References

23. Gagnon, J.A. et al. Efficient mutagenesis by Cas9 protein-mediated oligonucleotide insertion and large-scale assessment of single-guide RNAs. PLoS One 9, e98186 (2014).

24. Clement, K. et al. CRISPResso2 provides accurate and rapid genome editing sequence analysis. Nat Biotechnol 37, 224–226 (2019).

